# Metaphloem development in the Arabidopsis root tip

**DOI:** 10.1101/2021.04.30.442082

**Authors:** Moritz Graeff, Christian S. Hardtke

## Abstract

The phloem transport network is a major evolutionary innovation that enabled plants to dominate terrestrial ecosystems. In the growth apices, the meristems, apical stem cells continuously produce early, so-called protophloem. This is easily observed in Arabidopsis root meristems, where the differentiation of individual protophloem sieve element precursors into interconnected, conducting sieve tubes is laid out in a spatio-temporal gradient. The mature protophloem eventually collapses as the neighboring metaphloem takes over its function further distal from the stem cell niche. Compared to protophloem, metaphloem ontogenesis is poorly characterized, primarily because its visualization is challenging. Here we describe an improved protocol to investigate metaphloem development in Arabidopsis root tips in combination with a set of new molecular markers. We found that mature metaphloem sieve elements are only observed in the late post-meristematic root although their specification is initiated as soon as protophloem sieve elements enucleate. Moreover, unlike protophloem sieve elements, metaphloem sieve elements only differentiate once they have fully elongated. Finally, our results suggest that metaphloem differentiation is not directly controlled by protophloem-derived cues but rather follows a distinct, robust developmental trajectory.

**Summary statement:** Metaphloem sieve element differentiation in Arabidopsis roots follows a robust developmental trajectory.

## INTRODUCTION

The evolution of vascular tissues enabled plants to conquer land because it allowed the separation of the sites of photosynthesis from the sites of nutrient and water acquisition (Lucas et al., 2013). In extant angiosperms, the xylem vessels form hollow tubes to transport water and inorganic ions from the root system to the shoot system. This transport is mainly driven by the water potential differential between the soil and the atmosphere, and therefore by purely physical forces (Endo et al., 2019; Pratt and Jacobsen, 2017). Closely associated with the xylem is the phloem, which is composed of inter-connected sieve elements that form the conducting sieve tubes and their neighboring companion cells. Unlike xylem vessels, sieve elements are not dead, but during their differentiation process, they drastically alter their cellular makeup to optimize the transport flow. Most noticeable, they lose their nucleus and vacuole. Thus, sieve elements depend on the neighboring companion cells for the maintenance of their transport functions. Phloem sieve tubes mediate the long distance bulk transport of phloem sap, a viscous mix of sugars, metabolites as well as systemic signaling molecules, from source to sink organs, for example from mature photosynthesizing leaves to roots (Lopez-Salmeron et al., 2019). This transport is driven by a differential in osmotic pressure, which builds up through the controlled loading of osmotic sugars in the source tissue phloem and their unloading in the sink tissue phloem (Knoblauch et al., 2016; Zhang and Turgeon, 2018). The growth apices of plants, the meristems, are terminal sinks, whose activity is sustained by phloem sap delivered through the early, so-called protophloem. In root meristems, protophloem is produced by apical stem cells that reside adjacent to the quiescent center (QC) and matures while neighboring tissues still divide or undergo expansion growth (Esau, 1977; Lopez-Salmeron et al., 2019). Eventually, its sieve elements become non-functional and are completely obliterated as protophloem is replaced by emerging metaphloem. Although the metaphloem sieve elements share a common precursor with protophloem sieve elements (Bonke et al., 2003; Rodriguez-Villalon et al., 2014), the metaphloem only matures after the expansion growth of the surrounding tissues is completed (Esau, 1977). Metaphloem is then retained as the main conducting phloem, although it can later be replaced by secondary phloem in species that undergo secondary growth.

Non-invasive investigation of phloem development is challenging, on the one hand because sieve elements are thin and highly anisotropic cells, and on the other hand because the phloem is buried deep inside plant organs. Routine observation of protophloem by confocal microscopy is however possible in the root tip of *Arabidopsis thaliana* (Arabidopsis), where its development is laid out in a spatio-temporal gradient of ~20 cells from stem cell daughter to mature sieve element (Furuta et al., 2014; Rodriguez-Villalon et al., 2014). Arabidopsis root tips produce two protophloem strands, which are arranged opposite each other inside the stele, flanking an axis of xylem cells (Fig. 1A). The last two decades have seen tremendous advances in our understanding of protophloem ontogeny. Through its dissection by genetic approaches, numerous protophloem-specific mutants and molecular markers have become available. These studies underline the essential character of root protophloem, whose absence or disturbed development has grave, systemic consequences on root meristem growth and maintenance (Anne and Hardtke, 2017; Bonke et al., 2003; Rodriguez-Villalon et al., 2014). Whether the defects in the protophloem of pertinent mutants also extend to metaphloem remains largely unknown, mainly because of the difficulty in visualizing metaphloem development and a paucity of specific molecular markers for non-invasive investigation. Here we set out to mend this gap by developing a toolbox for the analysis of metaphloem development.

**Fig. 1.**
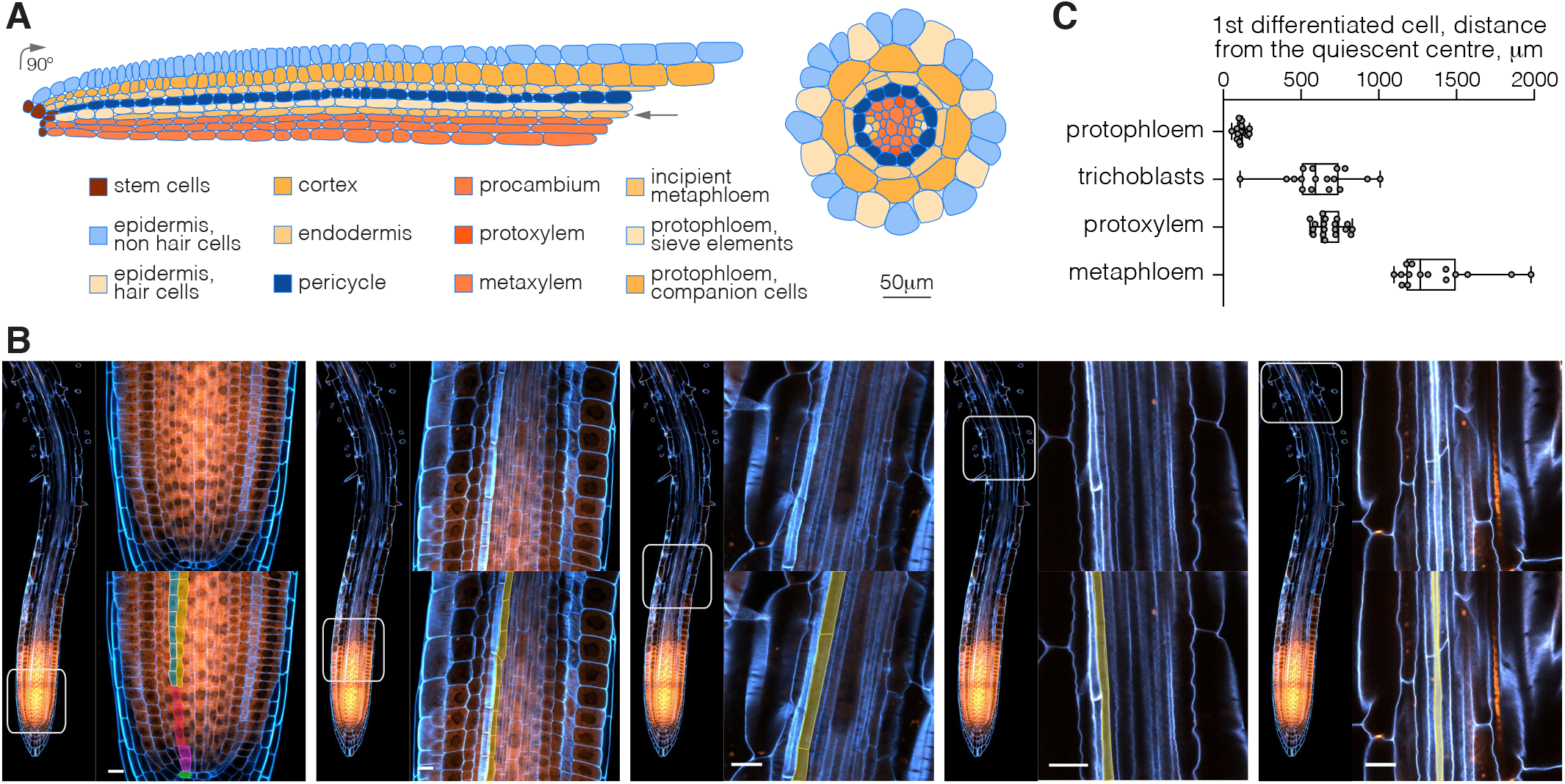
Development of metaphloem sieve elements (MPSEs) in the Arabidopsis root tip. (A) Schematic overview of tissue arrangement and development in an Arabidopsis root meristem, based on confocal microscopy images of a longitudinal half section and a horizontal cross section. (B) Confocal microscopy, optical sections illustrating MPSE development in a 7-day-old Arabidopsis Col-0 wildtype root tip stained with calcofluor white (CCFW; blue fluorescence) and propidium iodide (PI; reddish fluorescence) using the optimized “TetSee” protocol. Left overview panels indicate the approximate positions of the magnified images in the right panels. The bottom right panels are labeled counterparts of the raw images in the corresponding top right panels. The common stem cell precursors for the protophloem sieve element (PPSE) and MPSE cell files are labeled in red in the left-most assembly. Note the formative division giving rise to the developing PPSE strand (labeled green) and incipient MPSE strand (labeled yellow). Size bars are 10 micrometer. (C) Distance of the first visibly differentiated cell from the quiescent center, for different root tissues. Box plots display 2nd and 3rd quartiles and the median, bars indicate maximum and minimum.

## RESULTS AND DISCUSSION

### An optimized protocol for metaphloem visualization by confocal microscopy

Imaging of the Arabidopsis root tip by confocal microscopy techniques is routine but can be tricky depending on the tissue targeted for investigation. In particular, this applies to the cells inside the stele, which are small in diameter compared to the surrounding ground tissue or epidermis (Fig. 1A). For instance, whereas the diameter of cortex cells reaches ~25 to ~50 micrometers, protophloem cells are a mere ~5 micrometers across and therefore roughly 20 times smaller in their horizontal cross section profile (Fig. 1A). Together, the vascular tissues inside the stele occupy merely ~10% of the area in a root meristem cross section, although they represent ~40% of cell files. Despite progress in staining and fixation techniques, visualization of these cells can sometimes be challenging. For instance, while developing protophloem sieve elements (PPSEs) can be readily identified because of their early differentiation and associated cell wall build up (Truernit, 2014), mature, enucleated PPSEs are difficult to observe. Initially, PPSEs elongate from the ~20 micrometer typical of dividing cells to an intermediate stage of ~50 micrometer during which the principal differentiation steps occur. Once they are enucleated, they still elongate rapidly to about twice their length as they become the conductive unloading terminus of the PPSE cell file (Ross-Elliott et al., 2017). It is likely their high anisotropy in combination with a still elongating, soft cell wall that is responsible for the compression of maturing PPSEs by neighboring tissues once they lose their elevated turgor during fixation. Elongating cells possess relatively soft primary cell walls to facilitate directional expansion. Stabilizing secondary cell walls are only deposited during the final stages of differentiation, when the cells have reached their final size and adapt to their future roles. The phenomenon of cell shrinkage or collapse upon fixation is generally observed once all tissues have started to elongate further distal, in the generic cell elongation zone of the root meristem (Fig. S1A). The developing metaphloem sieve elements (MPSEs) are thus particularly affected, rendering their observation difficult with existing standard protocols, like chloral hydrate clearing (McBryde, 1936) or mPS-PI staining (Truernit et al., 2008). By contrast, the recently developed ClearSee (Kurihara et al., 2015) and TDE (2’2-thiodiethanol) clearing (Musielak et al., 2016) protocols not only preserved the structure of this delicate area (Fig. S1B) but also the fluorescence of reporter proteins.

Starting from these recent advances, we sought to develop a protocol that would leave the elongation area intact and permit routine observation of MPSEs. Through a test series with various combinations and concentrations of described detergents, clearing agents and fixation steps (Kurihara et al., 2015; Musielak et al., 2016; Ursache et al., 2018), we established an optimized procedure that maintained the integrity of the root elongation zone and allowed us to observe the progressive development of MPSE cell files (the “TetSee” protocol, see Materials and Methods) (Fig. S1C). Starting from the second formative division in the phloem lineage, the division that gives rise to the PPSE and MPSE cell files (Bonke et al., 2003; Rodriguez-Villalon et al., 2014), we could follow MPSE files across overlapping 3D renderings of serial confocal microscopy images (Fig. 1B). The morphologically visible onset of MPSE differentiation, as judged by intensified calcofluor white cell wall staining, was on average observed as far as ~1,400 micrometers from the QC. This was substantially later than the onset of morphological differentiation of PPSEs (~120 micrometer from the QC), trichoblasts (~620 micrometer from the QC) or protoxylem (~680 micrometer from the QC) (Fig. 1C). Thus, MPSEs only differentiated visibly once all other tissues had already matured, with the exception of the metaxylem, which differentiated around the same time or slightly later.

### Metaphloem sieve elements differentiate after they have reached their final cell size

Interestingly, whereas cell elongation and differentiation are tightly linked in PPSEs (Furuta et al., 2014; Rodriguez-Villalon et al., 2014), MPSEs elongated to roughly their final size before any cell wall build up became apparent (Fig. 1B). Observation of other cellular rearrangements indicative of MPSE differentiation, notably enucleation, proved to be difficult because of the high anisotropy of MPSEs and against the background from neighboring tissues, for example when nucleic acid dyes such as DAPI were used. However, our morphology-based observations were corroborated by analyses of a generic molecular marker of cellular differentiation in Arabidopsis, the MINIYO (IYO) protein (Sanmartin et al., 2011). Constitutively expressed IYO-GFP fusion protein is barely visible in the cytosol but accumulates in the nucleus once cells differentiate. In the root tip, IYO-GFP was therefore clearly visible in the quickly differentiating distal root tissues, the columella and lateral root cap (Fig. 2A). Among the proximal root tissues, protophloem is the first to differentiate and consistently, nuclear IYO-GFP accumulation became first apparent in differentiating PPSEs (Fig. 2A) (Sanmartin et al., 2011). Interestingly, they were followed by their companion cells with some distance, suggesting that PPSE companion cells only differentiate once PPSEs are fully elongated and functional (Fig. 2A). In the stele, developing protoxylem displayed nuclear IYO-GFP next (Fig. 2B), followed with some delay by MPSE cell files (Fig. 2C). In fact, nuclear IYO-GFP accumulation was only observed in developing MPSEs after developing protoxylem vessels had already completed their secondary wall build up and after they had themselves fully elongated (Fig. 2D). In summary, both our morphological and molecular analyses suggest that unlike in PPSEs, cell elongation and terminal differentiation do not coincide in MPSEs.

**Fig. 2.**
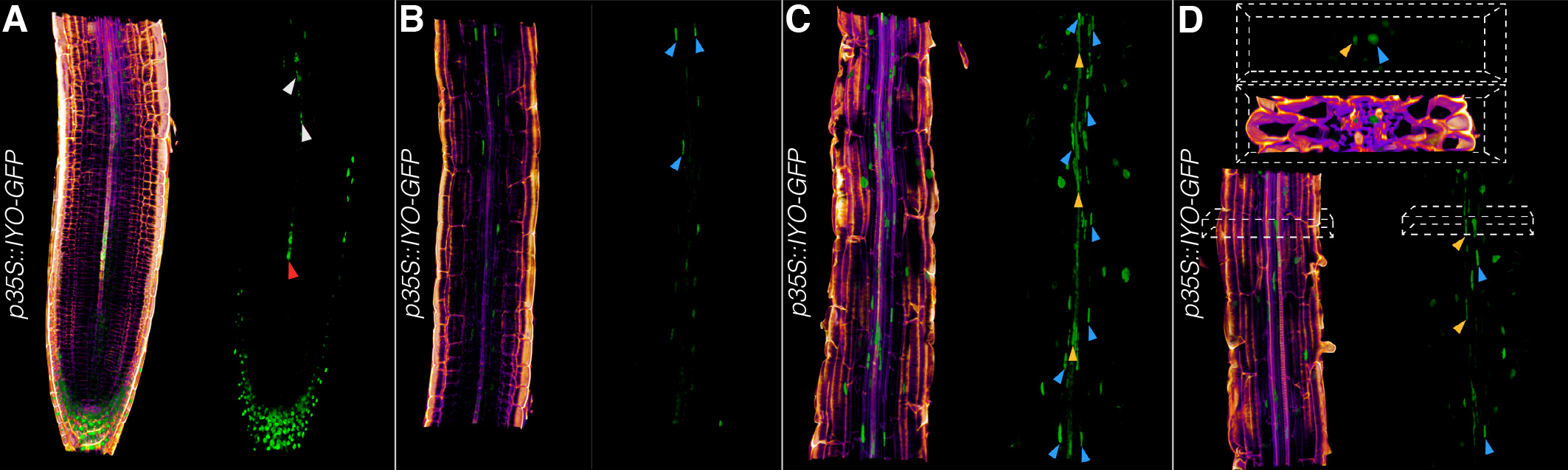
Differentiation timing in the Arabidopsis root tip. (A-D) 3D renderings of confocal image stacks, focused on the vasculature. Consecutive sections of a 7-day-old CCFW-stained root expressing the IYO-GFP fusion protein (green fluorescence) under control of the constitutive *35S* promoter are shown. Left panels: CCFW-GFP fluorescence composite images; right panels: GFP fluorescence only. Nuclear IYO-GFP accumulation indicates cellular differentiation. Differentiating PPSEs are pointed out by a red arrowhead, their companion cells by white arrowheads (A). Protoxylem cells (blue arrowheads) start to differentiate before MPSEs (orange arrowheads) (B), who enter differentiation once secondary wall build up in protoxylem cells becomes apparent (C,D).

### A set of new molecular markers for the investigation of metaphloem development

Although nuclear IYO accumulation is a very useful generic indicator of the onset of cellular differentiation (Sanmartin et al., 2011), it is not a marker for cell specification. We therefore sought to identify tissue-specific molecular markers that would allow us to trace the incipient beginnings of MPSE development. To this end, we mined the literature for genes that are specifically expressed in mature phloem in other contexts (Anstead et al., 2012; Bonke et al., 2003; Cayla et al., 2015; Khan et al., 2007; Sankar et al., 2014) and chose seven genes for further investigation. Moreover, we intersected existing phloem-related gene expression data sets (Brady et al., 2007; Clark et al., 2019; Kondo et al., 2016; Zhao et al., 2005) to identify a set of 14 additional metaphloem marker candidates. For some of them, existing reporter plasmids could be obtained, but for the majority we cloned promoter constructs that drive the expression of a nuclear localized fluorescent reporter (NLS-CITRINE). After their transformation into Col-0 wildtype plants, eight out of the 21 reporters showed activity in developing root phloem: the described *SISTER OF ALTERED PHLOEM DEVELOPMENT (SAPL)* (Ross-Elliott et al., 2017), *EARLY NODULIN-LIKE 9 (ENODL9)* (Khan et al., 2007), *SIEVE ELEMENT OCCLUSION-RELATED 2 (SEOR2)* (Anstead et al., 2012) and *SECONDARY WALL-ASSOCIATED NAC DOMAIN PROTEIN 2 (SND2)* (Kim et al., 2020) reporters (Fig. 3A-D); and the new reporters *DESIGUAL 2 (DEAL2)* (Wilson-Sanchez et al., 2018), *SIEVE ELEMENT MARKER 1* (*SEMA1*; AT2G35585), *SEMA2* (AT1G61760) and *SEMA3* (AT3G26350) (Fig. 4A-D).

**Fig. 3.**
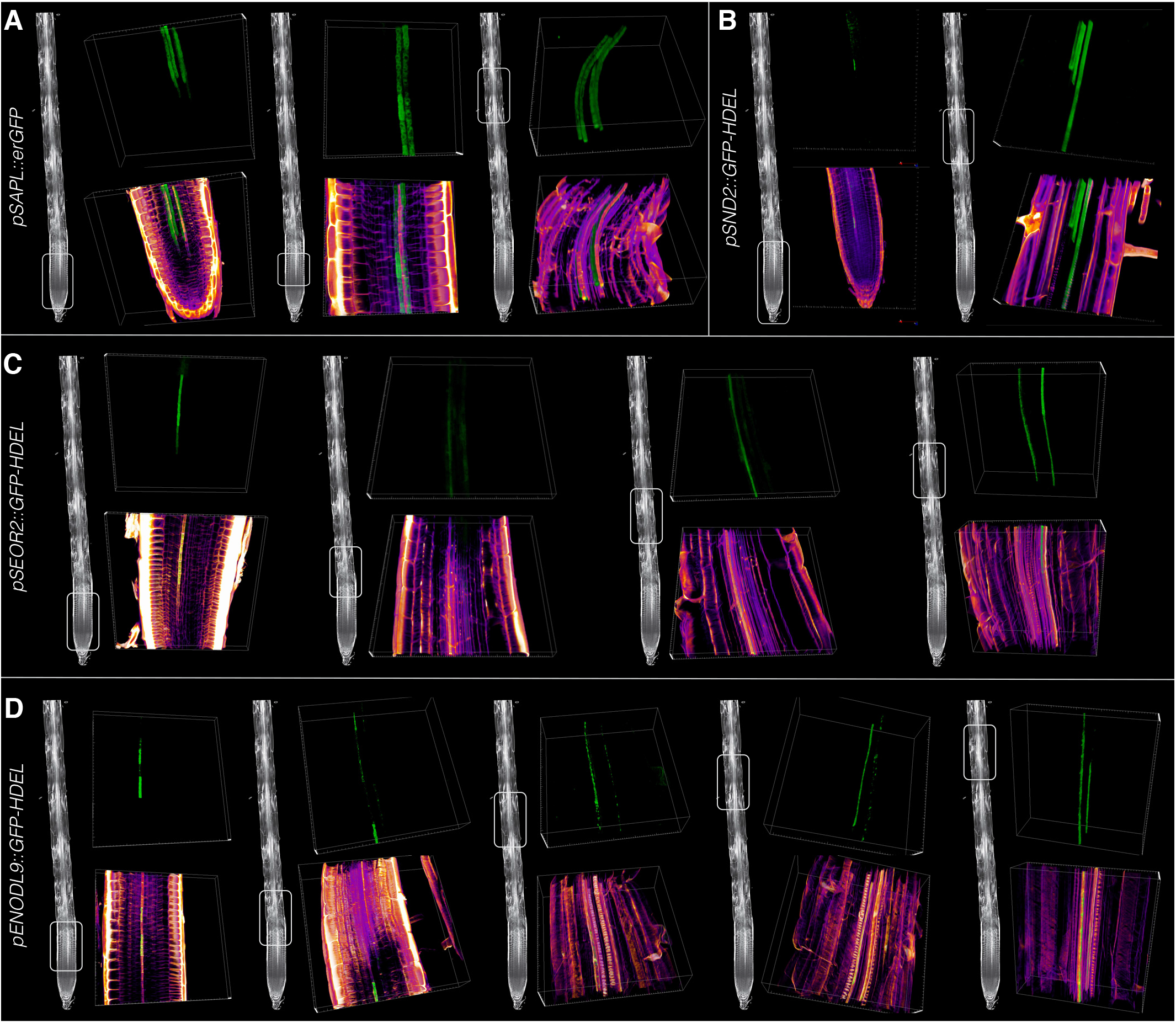
Reporter genes for phloem specification and differentiation I. (A-D) 3D renderings of confocal image stacks, focused on the vasculature of 7-day-old CCFW-stained roots that express the indicated reporter genes (green fluorescence). Left overview panels (generic wildtype root) indicate the approximate positions of the magnified images in the right panels. Bottom panels: CCFW-GFP fluorescence composite images; top panels: GFP fluorescence only. Note that for better visibility of details, images are not always to the same scale.

**Fig. 4.**
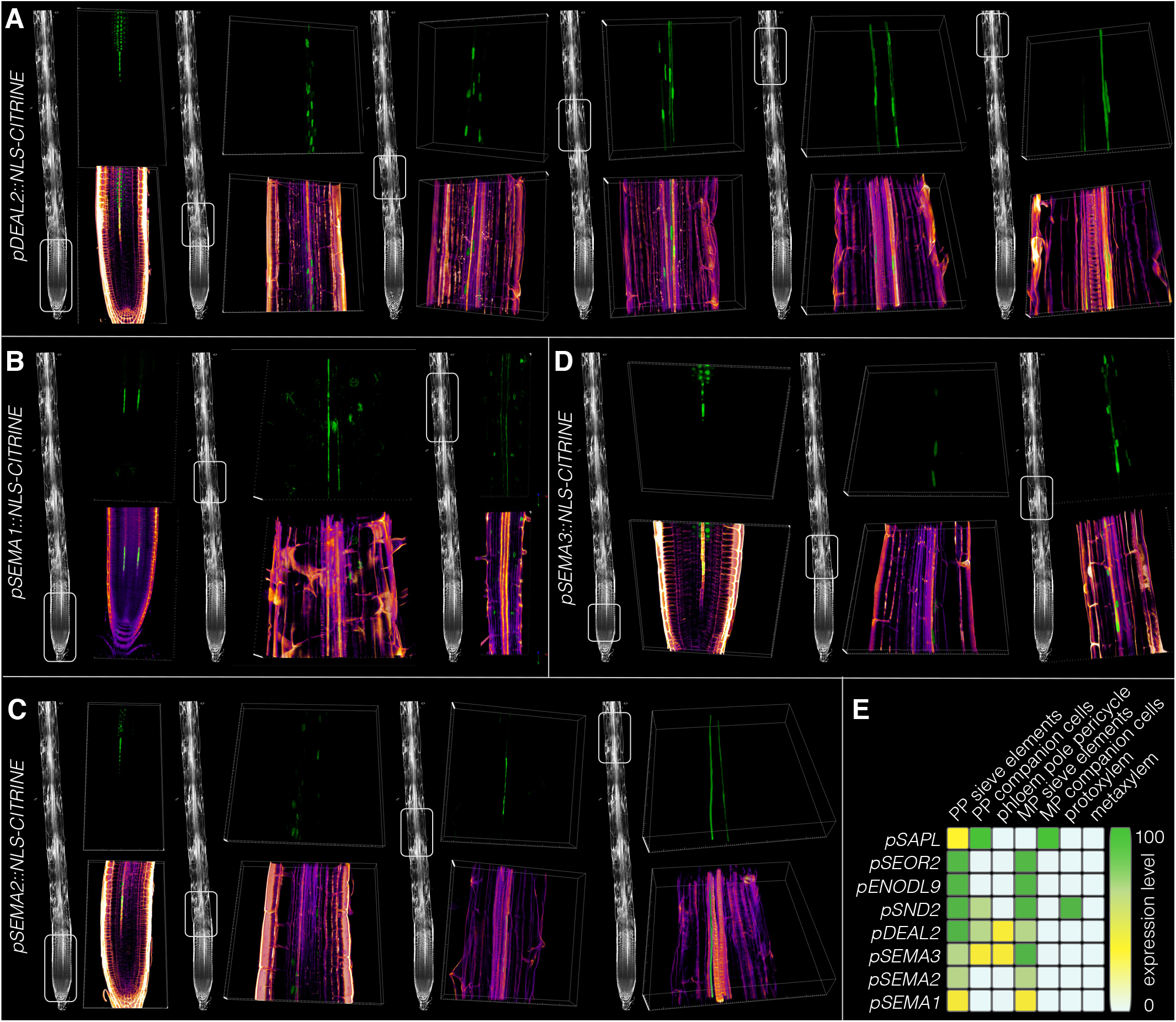
Reporter genes for phloem specification and differentiation II. (A-D) 3D renderings of confocal image stacks, focused on the vasculature of 7-day-old CCFW-stained roots that express the indicated reporter genes (green fluorescence). Left overview panels (generic wildtype root) indicate the approximate positions of the magnified images in the right panels. Bottom panels: CCFW-GFP fluorescence composite images; top panels: GFP fluorescence only. (E) Schematic summary of the tissue-specific expression patterns for the reporter genes shown in figures 3 and 4.

However, none of the reporters was exclusively active in the (incipient) metaphloem, rather, all markers were also expressed in the late developing protophloem. Among them, the *SAPL* expression was particular, because although it was expressed in late differentiating PPSEs similar to the other markers, thereafter it was highly specific for companion cells, both in the proto- and metaphloem, and not detected in developing MPSEs (Fig. 3A). Notably, *SAPL* was continuously expressed from the early coincidence with PPSE differentiation onward beyond differentiated MPSEs and was not observed in any other cell file. This suggests that the four companion cell files subsequently serve both PPSE and MPSE maintenance. The other markers were expressed in PPSEs as well as MPSEs, with varying levels of specificity. All of them were expressed in developing PPSEs, after the onset of cell wall build up and coincident with the partial elongation that occurs before enucleation. *ENODL9*, *SEOR2* and *SEMA1* were most specific for developing sieve elements (Figs. 3C,D and 4B). However, whereas *SEOR2* and *SEMA1* expression gradually ceased upon PPSE differentiation and only became active again later, *ENODL9* expression switched to the incipient MPSE file earlier and stayed on until MPSE differentiation terminated (Fig. 3D). Moreover, *SEOR2* expression reappeared earlier than *SEMA1* expression (Figs. 3C and 4B). The other reporters also displayed some marked expression outside of PPSEs/MPSEs. The *SND2* reporter was strongly expressed in late developing MPSEs, however it was also observed in developing metaxylem (Fig. 3B). *DEAL2*, *SEMA2* and *SEMA3* all switched expression to the cell files surrounding PPSEs after enucleation (Fig. 4A,C,D). In the metaphloem, *DEAL2* was expressed in MPSEs but also in the directly neighboring cell files, likely the companion cells (Fig. 4A). A similar pattern was observed for *SEMA3* (Fig. 4D), whereas *SEMA2* appeared to be specific for MPSEs (Fig. 4C). In summary, we were able to identify a set of reporters for metaphloem development that mark different stages as well as cell types (Fig. 4E). Their investigation confirmed that unlike what has been reported for PPSEs, cell elongation and differentiation are uncoupled in MPSEs, and also show that both sieve element types are associated with the same companion cell files.

### Metaphloem development is not affected by CLE45 treatment

The continuous expression of *ENODL9* in the MPSE cell files as soon as PPSEs enucleate also suggested that MPSE specification starts as soon as PPSE development is finished. This could mean that premature MPSE differentiation is prevented by lateral inhibition through cues derived from developing PPSEs. One such candidate signal are secreted CLAVATA3/EMBRYO SURROUNDING REGION-RELATED (CLE) signaling peptides, because low concentrations of certain synthetic CLE peptides suppress PPSE development when applied to roots (Depuydt et al., 2013; Hazak et al., 2017; Ito et al., 2006; Kinoshita et al., 2007; Rodriguez-Villalon et al., 2014), as does dosage increase of CLE45 (Czyzewicz et al., 2015b; Rodriguez-Villalon et al., 2014). Interestingly, CLE45 as well as CLE26 and CLE25 are specifically expressed in developing PPSEs (Czyzewicz et al., 2015a; Ren et al., 2019; Rodriguez-Villalon et al., 2014; Rodriguez-Villalon et al., 2015). CLE peptide signaling is however apparently not strictly required for protophloem development (Anne et al., 2018; Fukuda and Hardtke, 2020), rather it appears to act as a safeguard mechanism that maintains plasticity of phloem pole cells during their meristematic stage (Gujas et al., 2020).

Upon CLE45 treatment, the expression of both markers tested, *SEMA3* and *SEOR2*, disappeared from the protophloem, consistent with their prohibitive effect on PPSE formation (Fig. 5A-D). However, both markers persisted in developing MPSEs (Fig. 5B,D), in line with the observation that their differentiation appeared unaffected. Notably, this observation also confirmed once more that MPSE specification is position-rather than lineage-dependent, because the PPSE and MPSE cell files arise from the same stem cell daughter through a periclinal division that is suppressed by CLE45 application (Rodriguez-Villalon et al., 2014; Rodriguez-Villalon et al., 2015). Moreover, the absence of pertinent phenotypes in *cle25* mutants (Ren et al., 2019) as well as in receptor mutants that are fully insensitive against all three CLE peptides (Anne et al., 2018) corroborates the conclusion that PPSE-derived CLE peptides do not impinge on MPSE development under normal circumstances. In summary, CLE45 peptide treatment efficiently suppressed PPSE formation, but did not interfere with MPSE development.

**Fig. 5.**
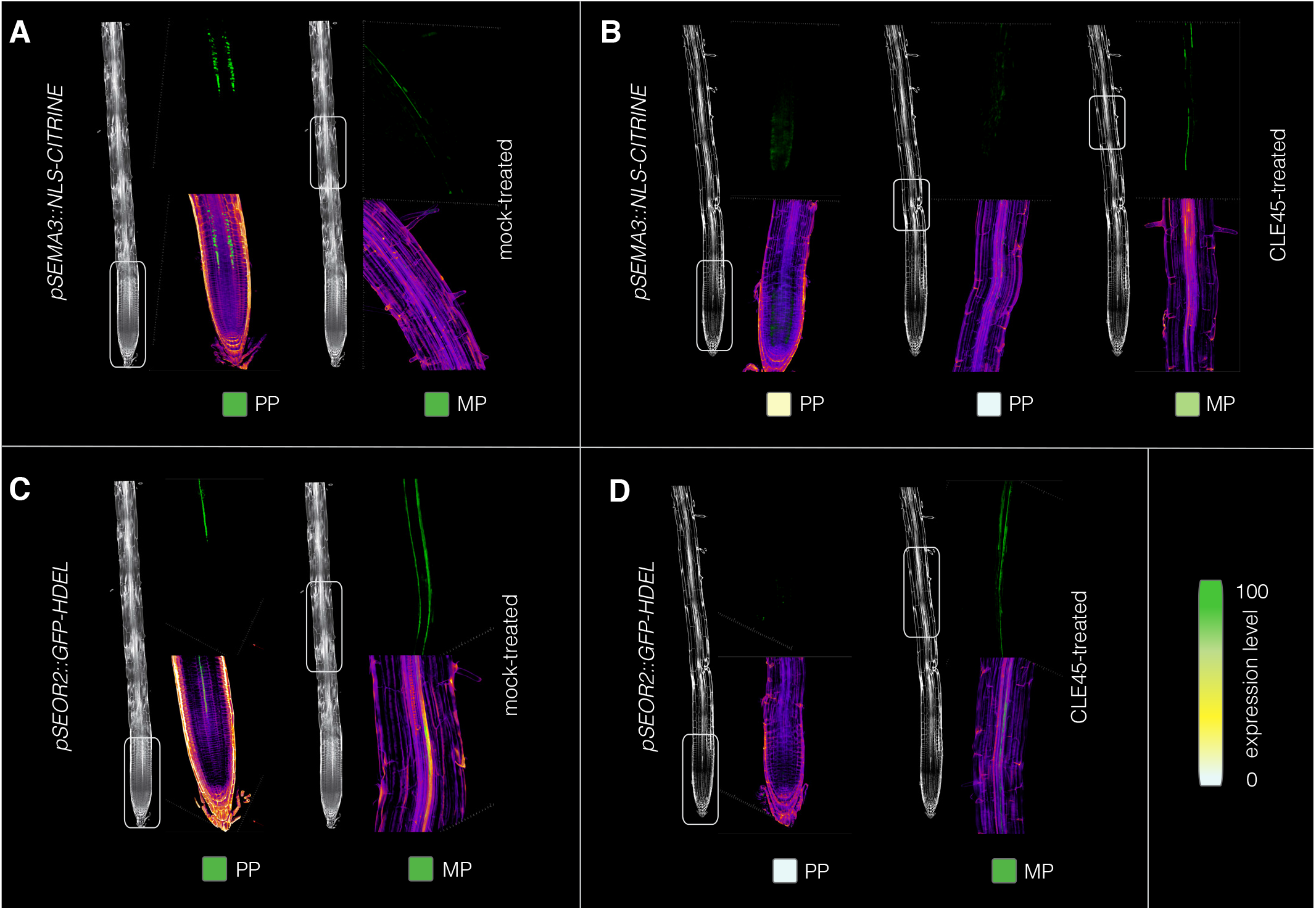
MPSE reporter genes do not respond to CLE45-treatment. (A-D) 3D renderings of confocal image stacks, focused on the vasculature of 7-day-old CCFW-stained roots that express the indicated reporter genes (green fluorescence). Left overview panels (generic wildtype root, treated with mock or CLE45) indicate the approximate positions of the magnified images in the right panels. Bottom panels: CCFW-GFP fluorescence composite images; top panels: GFP fluorescence only. Roots were either grown on mock (A,C) or 15 nM CLE45 peptide (B,D).

### MPSE development follows a robust developmental trajectory

In the protophloem, CLE45 signaling is quantitatively antagonized by the vascular plant -specific *OCTOPUS (OPS)* gene. *OPS* is thus a positive regulator of PPSE differentiation that is expressed from early on in the protophloem and insulates developing PPSEs against the effects of autocrine CLE45 signalling (Breda et al., 2017; Breda et al., 2019). In *ops* loss-of-function mutants, developing PPSEs frequently fail to differentiate (including failure to build up cell wall and thereby appearing as so-called gap cells), which causes discontinuities in the protophloem strands and disturbs the transport of phloem sap into the meristem (Anne and Hardtke, 2017; Rodriguez-Villalon et al., 2014; Truernit et al., 2012). *OPS* is also weakly expressed in the incipient MPSE cell file, against a background of low, ubiquitous expression of its homolog *OPS-LIKE 2 (OPL2)* that increases in developing metaphloem (Ruiz Sola et al., 2017). Whereas *opl2* single mutants did not display apparent phenotypes, except (in our hands) a more variable root growth vigor (Fig. S2A), the *ops opl2* double mutant is the only described genotype with MPSE defects so far (Fig. S2B) (Ruiz Sola et al., 2017), apart from mutants that lack protophloem and metaphloem altogether. Compared to *ops* single mutants, root growth vigor was further diminished in *ops opl2* double mutants (Fig. S2A,C) and they also displayed aggravated PPSE differentiation defects (Fig. S2B,D) (Ruiz Sola et al., 2017). The latter were more severe than evident from simple gap cell presence-absence counts, because *ops opl2* double mutants often had only one distinguishable PPSE strand. To better understand how MPSE and PPSE differentiation is affected in *ops* single and *ops opl2* double mutants, we introduced some of our reporter genes into these backgrounds.

In *ops* single mutants, the *SEMA3* reporter was expressed at the later stages of PPSE differentiation as in wildtype, but absent in developing PPSEs that failed to differentiate (Fig. S3A), underlining their different cellular identity. In developing MPSEs, *SEMA3* expression appeared to be unaffected (Fig. S3B,C). By contrast, *SAPL* reporter activity was still observed in gap cells (Fig. S3D), which corroborates earlier observations and is in line with the recent proposal that they adopt companion cell identity (Gujas et al., 2020) as well as the strong continuous companion cell-specific *SAPL* expression after PPSE differentiation. Again, *SAPL* expression appeared unaffected in the developing metaphloem region of *ops* mutants (Fig. S3E). Together, these findings reiterate that the defects in *ops* mutants are protophloem-specific (Ruiz Sola et al., 2017; Truernit et al., 2012). Interestingly, *SAPL* expression could still be detected in protophloem gap cells later on (Fig. S3F), indicating that PPSE cells that fail to differentiate properly in the protophloem differentiation window fail to catch up.

In *ops opl2* double mutants, the *SEOR2*, *DEAL2*, *SEMA2* and *SEMA3* markers displayed normal expression, except their apparent absence in gap cells (Fig. 6A-D). Despite the described MPSE differentiation defects (Ruiz Sola et al., 2017), which we could also observe in optical cross sections (Fig. S2B), our markers were however essentially continuously expressed in developing MPSEs of *ops opl2* mutants (Fig. 6A-D). Thus, we could not detect corresponding “metaphloem gap cells”, possibly because differentiating MPSEs are quite long (200-300 micrometer) and because surveying extended stretches of MPSEs was difficult. Nevertheless, although *ops opl2* double mutants typically displayed marker expression in both PPSE and MPSE strands (Fig. 7A), this pattern also often deviated from wildtype. Upon closer inspection, this could be attributed to the reappearance of reporter expression in undifferentiated protophloem cell files long after the zone of normal PPSE differentiation, clearly visible from elongated PPSEs that expressed the respective marker (Fig. 7B). Thus, the observation that one of the two protophloem poles in *ops opl2* mutants was frequently absent (Fig. S2B) (Ruiz Sola et al., 2017) could also reflect a strongly delayed differentiation of one PPSE strand. *In extremis*, the delay was such that it overlapped with reporter expression in the neighboring MPSE cell files (Fig. 7C). We had not observed such atypical differentiation in *ops* single mutants. This not only indicates that PPSE differentiation can be substantially delayed in *ops opl2* mutants, but also that the onset of MPSE differentiation is largely independent of such delays. Corroborating the independent trajectory of MPSE differentiation, in *ops opl2* roots where only one PPSE cell file showed differentiating cells (any gap cells notwithstanding) and marker expression was also absent from the failed PPSE cell file later on, marker expression in both MPSE cell files appeared to be normal (Fig. 7D) and could typically be observed shortly after protoxylem cells with secondary cell walls were visible, as in wildtype. Together with the observed activity of our markers in CLE45-treated roots, our analyses therefore suggest that MPSE development follows a robust trajectory that is largely independent from PPSE development.

**Fig. 6.**
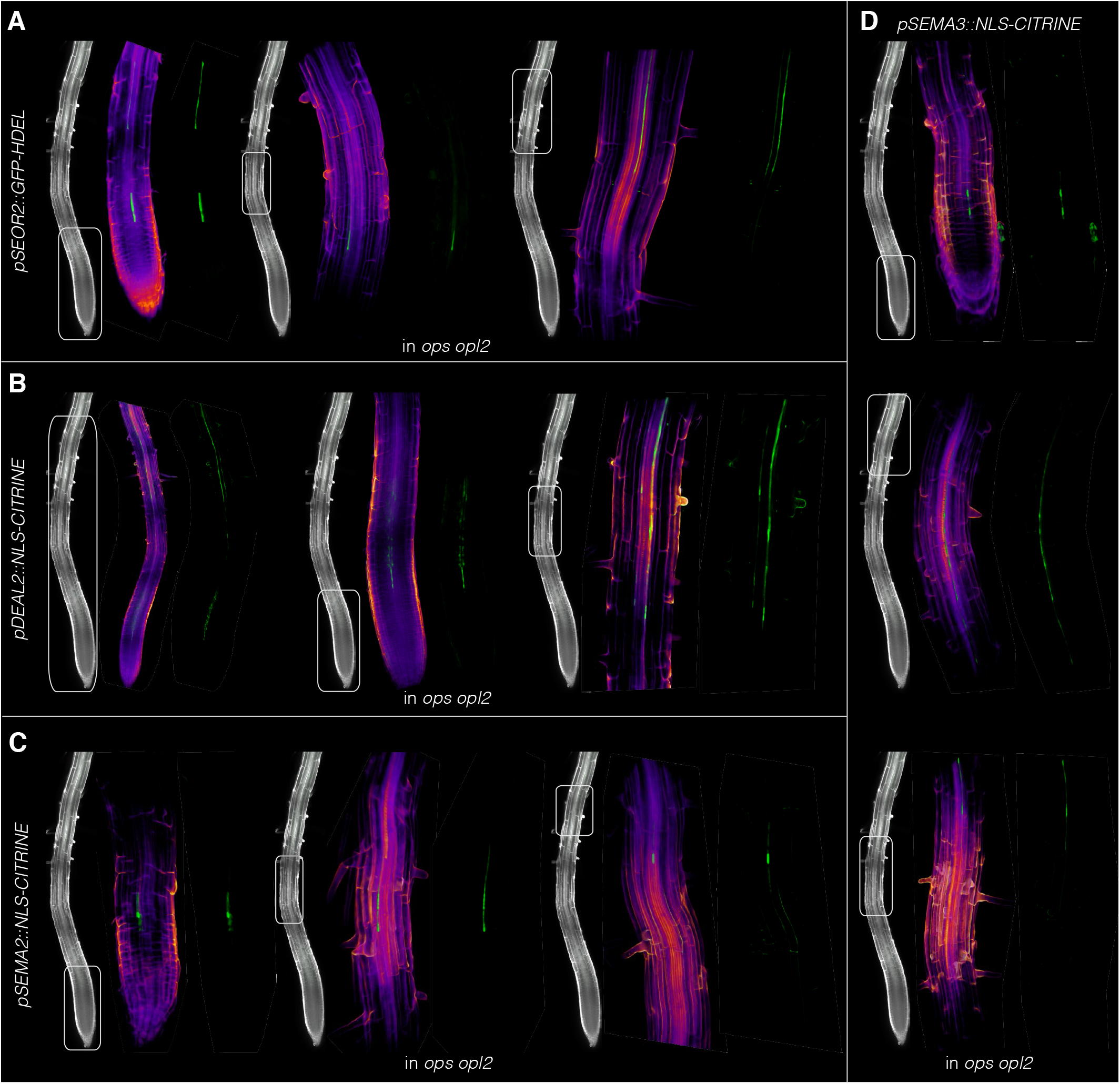
Phloem reporter gene expression in *ops opl2* double mutants I. (A-D) 3D renderings of confocal image stacks, focused on the vasculature of 7-day-old CCFW-stained roots that express the indicated reporter genes (green fluorescence) in *ops opl2* double mutant background. Left overview panels (generic *ops opl2* root) indicate the approximate positions of the magnified images in the right panels. Center panels: CCFW-GFP fluorescence composite images; right panels: GFP fluorescence only. Note that for better viewing of details, images are not always to the same scale.

**Fig. 7.**
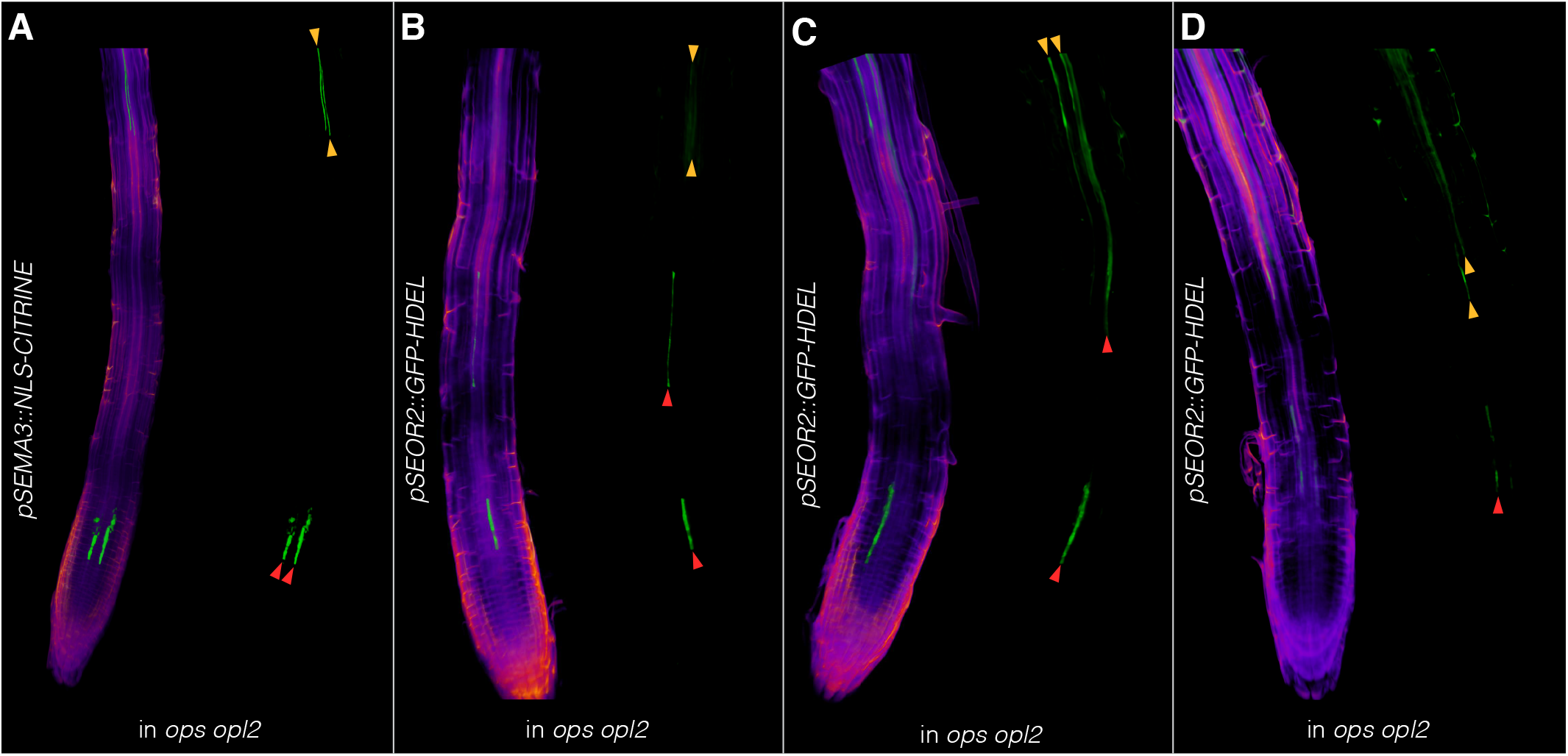
Phloem reporter gene expression in *ops opl2* double mutants II. (A-D) 3D renderings of confocal image stacks, focused on the vasculature of 7-day-old CCFW-stained roots that express the indicated reporter genes (green fluorescence) in *ops opl2* double mutant background. Left panels: CCFW-GFP fluorescence composite images; right panels: GFP fluorescence only. PPSE or MPSE cell files expressing molecular markers are pointed out by red or orange arrowheads, respectively.

Finally, it is noteworthy that OPS action is exquisitely dosage-sensitive (Breda et al., 2017; Breda et al., 2019) and promotes PPSE differentiation by quantitatively antagonizing CLE45 signaling via the receptor kinase BARELY ANY MERISTEM 3 (Breda et al., 2017; Breda et al., 2019; Fukuda and Hardtke, 2020). Moreover, an excess of ectopic OPS activity leads to premature differentiation across root tissues (Breda et al., 2019). Thus, our results are consistent with the notion that the differential expression levels of OPS family proteins such as *OPS* and *OPL2* along the gradient of developing PPSE versus MPSE cell files contributes to the correct spatio-temporal separation of their differentiation.

### A toolbox for the investigation of metaphloem development

In summary, our study extends our toolbox for the investigation of sieve element development in the Arabidopsis root, with a special focus on the so far poorly described differentiation of the metaphloem, and provides first forays into its genetic control. Our results highlight commonalities between PPSE and MPSE development, but also suggest that metaphloem development follows a robust trajectory that is not directly influenced by adjacent or preceding PPSE development under normal circumstances. Combined with state-of-the-art technical advances, such as single cell RNA sequencing or tissue-specific gene knock-out (Smetana et al., 2019; Wendrich et al., 2020), our observations should enable more targeted future approaches to dissect metaphloem development and discover its unique features.

## MATERIALS AND METHODS

Plant culture, transformation and common molecular biology followed previously described standard procedures (Cattaneo et al., 2019; Graeff et al., 2020; Kang et al., 2017).

### Plant materials and growth conditions

Seeds were surface sterilized using 3% sodium hypochlorite, sown onto half strength Murashige & Skoog agar medium (0.9% agarose) supplemented with 0.3% sucrose and stratified for 3 days at 4°C. Plants were grown under continuous white light (intensity ~120 μE) at 22°C. All mutants and marker lines were in the *Arabidopsis thaliana* Columbia-0 (Col-0) wild type background. The *ops* and *ops opl2* mutant lines were described previously (Ruiz Sola et al., 2017; Truernit et al., 2012). CLE45 peptide treatments were performed as described (Anne et al., 2018).

### Database mining and selection of sieve element marker candidates

For the selection of the sieve element (SE) marker candidates, expression of 576 genes enriched in cells expressing the S32 phloem marker (AT2G18380) was analysed along the root, in cells expressing *SUC2* (AT1G22710) (Brady et al., 2007), in cells expressing *CVP2* (AT1G05470) (Clark et al., 2019), in a general root and seedling gene expression dataset (Gan et al., 2011), and in the “VISUAL” phloem and xylem datasets (Kondo et al., 2016). Twenty candidate genes that i) showed expression in the phloem poles, ii) showed increased expression further away from the meristem, iii) showed relatively higher expression in the root, and iv) appeared in the VISUAL phloem datasets were tested as SE markers.

### Transgene constructs

For the construction of SE markers, respective promoter fragments of 1,500 bp to 2,500 bp were amplified from genomic Col-0 DNA using suitable oligonucleotides with overhangs (attB1/2 extensions for *ENODL9*, *SEOR2*, *SND2*, and attB4/1r extensions for *DEAL2* and *SEMA1-3*) for subsequent *Gateway™* cloning (see Table S1). The manufacturer’s solutions (*ThermoFisher Scientific* article numbers 11791020 and 11789020) and protocols were used for all cloning reactions. Amplified fragments were cloned into suitable entry vectors and the *ENODL9*, *SEOR2* and *SND2* promoters were transferred into the pMDC205 destination vector in front of a GFP reporter with an ER retention signal (Curtis and Grossniklaus, 2003). The *DEAL2* and *SEMA1-3* promoters were recombined together with NLS-CITRINE in a multisite gateway reaction into the pK7m24 vector backbone. Flowering Col-0 plants were transformed using the floral dip method and transformants were selected either on ½ MS media containing 25 mg/ml hygromycin B or 25 mg/ml kanamycin following a fast selection procedure (Harrison et al., 2006).

### Tissue fixation and clearing (TetSee protocol)

For microscopy, 7-day-old seedlings were fixed in a solution of 4% PFA in PBS buffer and transferred into a vacuum of 25 to 30 mmHg/Torr for 15 to 30 min. Subsequently, seedlings were washed three times in PBS for 5 min. For clearing of the samples, different protocols were used and assessed for the quality of the tissue preservation along the root. While standard protocols like chloral hydrate clearing (McBryde, 1936) or mPS-PI staining (Truernit et al., 2008) caused the shrinking of the cells in the early elongation zone, the recently developed ClearSee (Kurihara et al., 2015) and 2’2-thiodiethanol-clearing (Musielak et al., 2016) protocols preserved the structure of this delicate area as well as the fluorescence of reporter proteins. The two protocols were further optimized and combined into the “TetSee” (2’2-**T**hiodi**et**hanol-Clear**See**) protocol. Briefly, the washed seedlings were transferred into TetSee X solution (15% Na-deoxycholate, 25% urea, 10 % glycerol, 5% 2’2-thiodiethanol [*Merck*, Product No. 166782], 1% Triton X-100) and kept for 3 days at 4°C with daily changes of the TetSee X solution. For microscopy, the TetSee X solution was removed and replaced by TetSee solution (the TetSee X solution without Triton X-100) containing 0.25 mg/ml calcofluor white (CCFW; *Sigma*, Product No. F3543). Seedlings were incubated in the CCFW staining solution for 6 h or overnight, washed once in TetSee solution and then transferred onto microscopy slides with TetSee solution as mounting medium.

### Microscopy

For morphological assessment of phloem differentiation, roots were prepared as described above and the CCFW-stained cell walls were imaged using a *Zeiss* LSM880 confocal microscope with a 40x objective. A 405 nm laser was used for CCFW excitation, and the cell wall signal was recorded in a range from 450 to 480 nm. For imaging of the SE marker lines, GFP or CITRINE were sequentially excited with 488 nm and their emission recorded from 500 to 560 nm. Tile scans and Z scans were combined in order to obtain continuous images of the vasculature from the root meristem to the differentiated metaphloem. Additionally, a *Nikon* Spinning disc CSU-W1 confocal microscope with a 40x objective was used to record images of the CLE45-treated and the *ops opl2* SE marker lines. Analysis of the images and generation of 3D renderings from the Z stacks were performed using the *GNU icy* software.

## ACKNOWLEDGEMENTS

We would like to thank Dr. Y. Helariutta for a gift of the *pSAPL::erGFP* reporter plasmid, and Drs. J. Sanchez-Serrano and E. Rojo for a gift of the *p35S::IYO-GFP* plasmid and transgenic line.

## SUPPLEMENTARY FIGURE LEGENDS

**Fig. S1.**
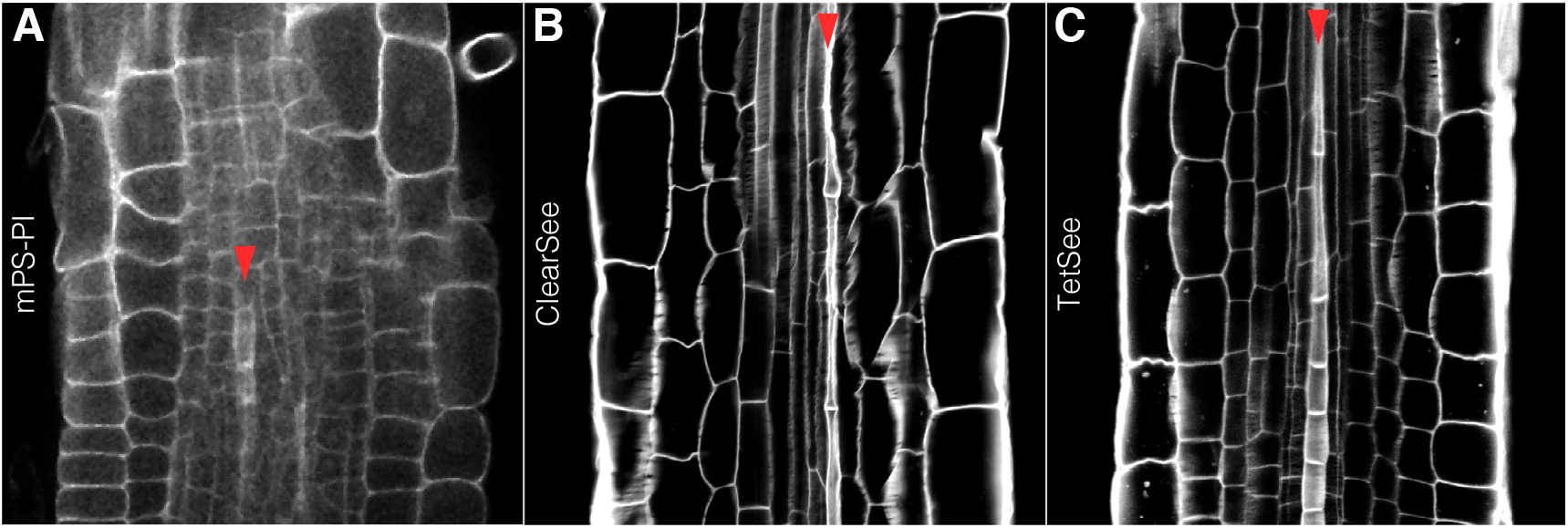
Imaging of elongating root cells using different fixation protocols. (A-C) Confocal microscopy, optical sections of 7-day-old CCFW-stained root meristems (white fluorescence) fixed with different protocols as indicated. Red arrowheads point out elongating PPSEs.

**Fig. S2.**
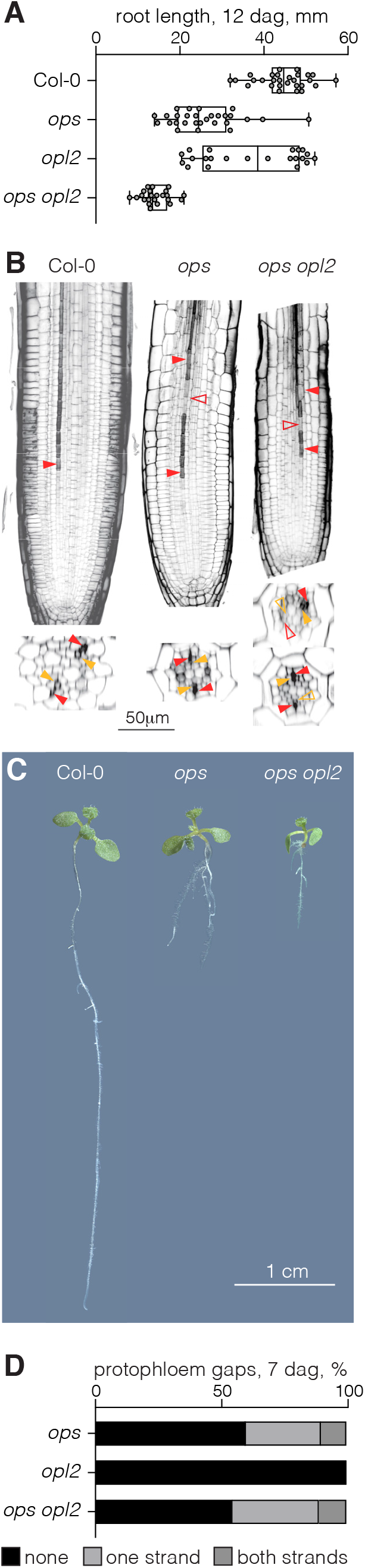
Root phenotypes of *ops opl2* double mutants. (A) Primary root length of 12-day-old seedlings of indicated genotypes. Box plots display 2nd and 3rd quartiles and the median, bars indicate maximum and minimum. (B) Confocal microscopy, longitudinal and horizontal optical cross sections of 7-day-old CCFW-stained root meristems (black fluorescence). Full red arrowheads point out normally differentiating PPSEs, open red arrowheads point out cells in the PPSE file that fail to differentiate (“gap cells). (C) Images of 12-day-old seedlings of the indicated genotypes. (D) Frequency of PPSE cell files with gap cells in root meristems of the indicated genotypes (n=20-30).

**Fig. S3.**
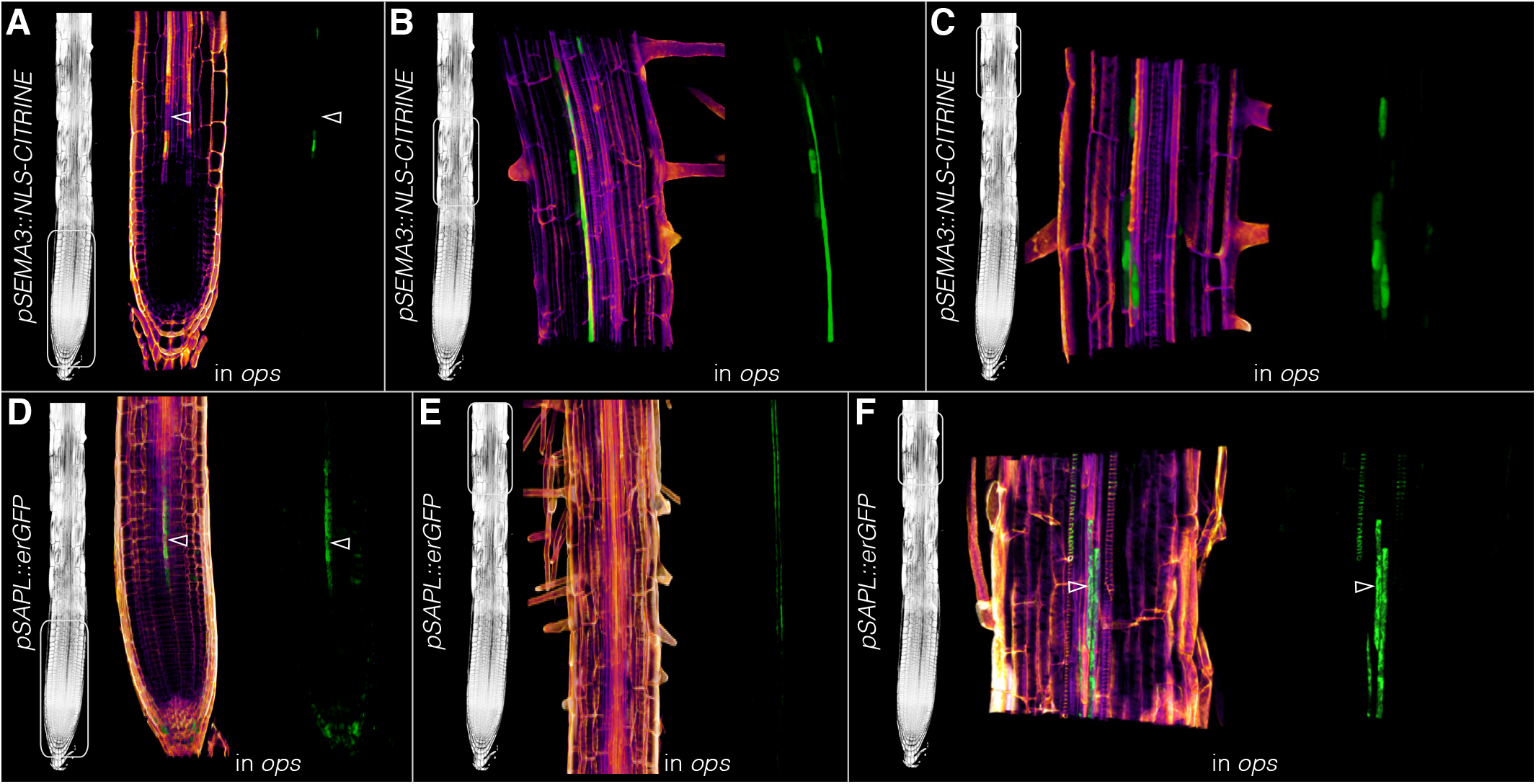
Phloem reporter gene expression in *ops* single mutants. (A-F) 3D renderings of confocal image stacks, focused on the vasculature of 7-day-old CCFW-stained roots that express the indicated reporter genes (green fluorescence) in *ops* single mutant background. Left overview panels (generic *ops* root) indicate the approximate positions of the magnified images in the right panels. Center panels: CCFW-GFP fluorescence composite images; top panels: GFP fluorescence only. Note that for better viewing of details, images are not always to the same scale. Open arrowheads in (A), (D) and (F) point out cells in the PPSE file that fail to differentiate (“gap cells).

---

**Table S1.**
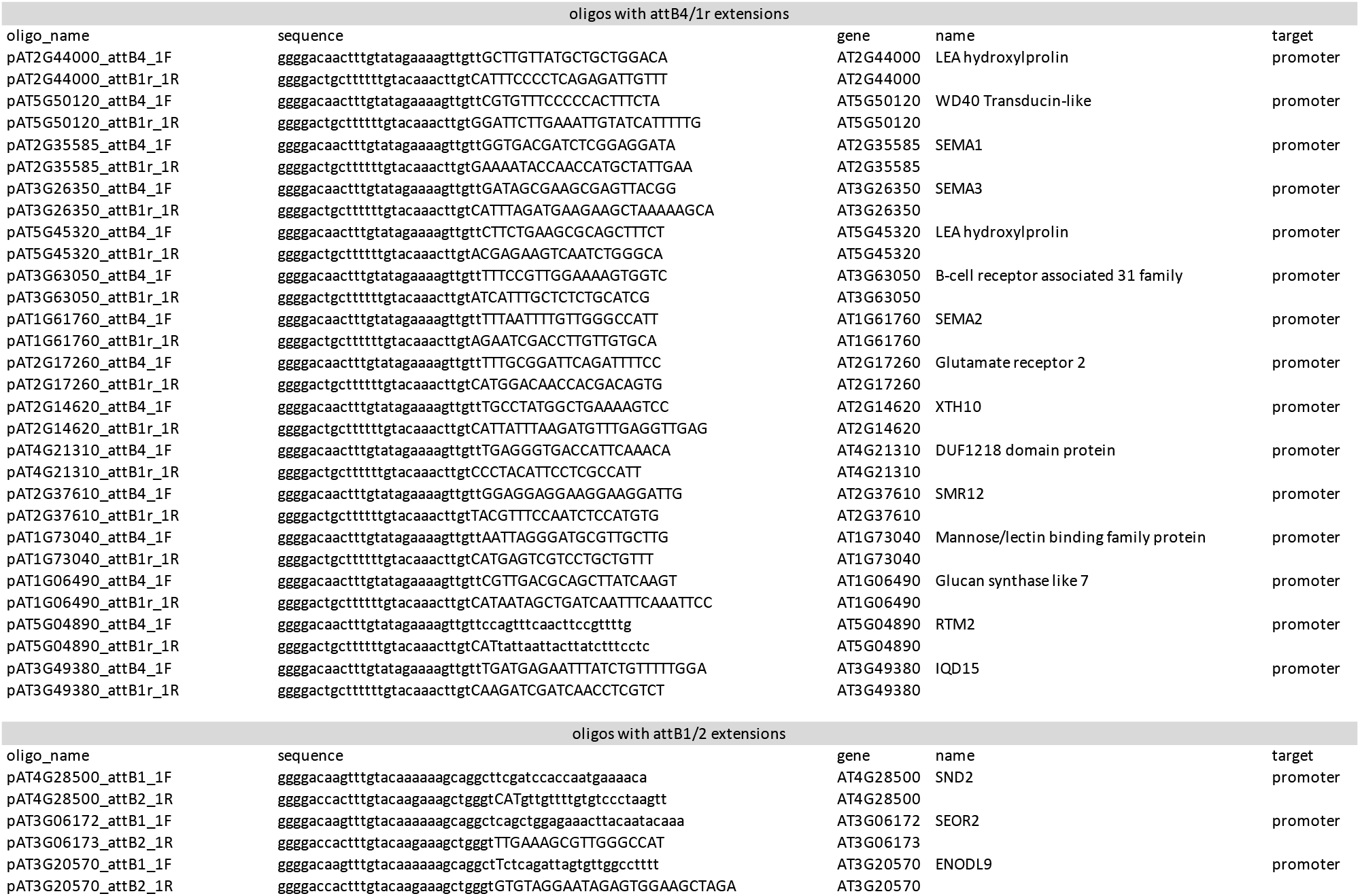
5’ to 3’ oligonucleotide sequences

## REFERENCES

Anne, P., Amiguet-Vercher, A., Brandt, B., Kalmbach, L., Geldner, N., Hothorn, M. and Hardtke, C. S. (2018). CLERK is a novel receptor kinase required for sensing of root-active CLE peptides in Arabidopsis. Development 145.

Anne, P. and Hardtke, C. S. (2017). Phloem function and development-biophysics meets genetics. Curr Opin Plant Biol 43, 22–28.

Anstead, J. A., Froelich, D. R., Knoblauch, M. and Thompson, G. A. (2012). Arabidopsis P-protein filament formation requires both AtSEOR1 and AtSEOR2. Plant Cell Physiol 53, 1033–1042.

Bonke, M., Thitamadee, S., Mahonen, A. P., Hauser, M. T. and Helariutta, Y. (2003). APL regulates vascular tissue identity in Arabidopsis. Nature 426, 181–186.

Brady, S. M., Orlando, D. A., Lee, J. Y., Wang, J. Y., Koch, J., Dinneny, J. R., Mace, D., Ohler, U. and Benfey, P. N. (2007). A high-resolution root spatiotemporal map reveals dominant expression patterns. Science 318, 801–806.

Breda, A. S., Hazak, O. and Hardtke, C. S. (2017). Phosphosite charge rather than shootward localization determines OCTOPUS activity in root protophloem. Proc Natl Acad Sci U S A 114, E5721–E5730.

Breda, A. S., Hazak, O., Schultz, P., Anne, P., Graeff, M., Simon, R. and Hardtke, C. S. (2019). A Cellular Insulator against CLE45 Peptide Signaling. Current biology : CB 29, 2501–2508 e2503.

Cattaneo, P., Graeff, M., Marhava, P. and Hardtke, C. S. (2019). Conditional effects of the epigenetic regulator JUMONJI 14 in Arabidopsis root growth. Development 146.

Cayla, T., Batailler, B., Le Hir, R., Revers, F., Anstead, J. A., Thompson, G. A., Grandjean, O. and Dinant, S. (2015). Live imaging of companion cells and sieve elements in Arabidopsis leaves. PloS one 10, e0118122.

Clark, N. M., Buckner, E., Fisher, A. P., Nelson, E. C., Nguyen, T. T., Simmons, A. R., de Luis Balaguer, M. A., Butler-Smith, T., Sheldon, P. J., Bergmann, D. C., et al. (2019). Stem-cell-ubiquitous genes spatiotemporally coordinate division through regulation of stem-cell-specific gene networks. Nature communications 10, 5574.

Curtis, M. D. and Grossniklaus, U. (2003). A gateway cloning vector set for high-throughput functional analysis of genes in planta. Plant Physiol 133, 462–469.

Czyzewicz, N., Shi, C. L., Vu, L. D., Van De Cotte, B., Hodgman, C., Butenko, M. A. and De Smet, I. (2015a). Modulation of Arabidopsis and monocot root architecture by CLAVATA3/EMBRYO SURROUNDING REGION 26 peptide. J Exp Bot 66, 5229–5243.

Czyzewicz, N., Wildhagen, M., Cattaneo, P., Stahl, Y., Pinto, K. G., Aalen, R. B., Butenko, M. A., Simon, R., Hardtke, C. S. and De Smet, I. (2015b). Antagonistic peptide technology for functional dissection of CLE peptides revisited. J Exp Bot 66, 5367–5374.

Depuydt, S., Rodriguez-Villalon, A., Santuari, L., Wyser-Rmili, C., Ragni, L. and Hardtke, C. S. (2013). Suppression of Arabidopsis protophloem differentiation and root meristem growth by CLE45 requires the receptor-like kinase BAM3. Proc Natl Acad Sci U S A 110, 7074–7079.

Endo, S., Iwai, Y. and Fukuda, H. (2019). Cargo-dependent and cell wall-associated xylem transport in Arabidopsis. New Phytol 222, 159–170.

Esau, K. (1977). Anatomy of seed plants (2d edn). New York: Wiley.

Fukuda, H. and Hardtke, C. S. (2020). Peptide Signaling Pathways in Vascular Differentiation. Plant Physiol 182, 1636–1644.

Furuta, K. M., Yadav, S. R., Lehesranta, S., Belevich, I., Miyashima, S., Heo, J. O., Vaten, A., Lindgren, O., De Rybel, B., Van Isterdael, G., et al. (2014). Plant development. Arabidopsis NAC45/86 direct sieve element morphogenesis culminating in enucleation. Science 345, 933–937.

Gan, X., Stegle, O., Behr, J., Steffen, J. G., Drewe, P., Hildebrand, K. L., Lyngsoe, R., Schultheiss, S. J., Osborne, E. J., Sreedharan, V. T., et al. (2011). Multiple reference genomes and transcriptomes for Arabidopsis thaliana. Nature 477, 419–423.

Graeff, M., Rana, S., Marhava, P., Moret, B. and Hardtke, C. S. (2020). Local and Systemic Effects of Brassinosteroid Perception in Developing Phloem. Current biology : CB 30, 1626–1638 e1623.

Gujas, B., Kastanaki, E., Sturchler, A., Cruz, T. M. D., Ruiz-Sola, M. A., Dreos, R., Eicke, S., Truernit, E. and Rodriguez-Villalon, A. (2020). A Reservoir of Pluripotent Phloem Cells Safeguards the Linear Developmental Trajectory of Protophloem Sieve Elements. Current biology : CB 30, 755–766 e754.

Harrison, S. J., Mott, E. K., Parsley, K., Aspinall, S., Gray, J. C. and Cottage, A. (2006). A rapid and robust method of identifying transformed Arabidopsis thaliana seedlings following floral dip transformation. Plant Methods 2, 19.

Hazak, O., Brandt, B., Cattaneo, P., Santiago, J., Rodriguez-Villalon, A., Hothorn, M. and Hardtke, C. S. (2017). Perception of root-active CLE peptides requires CORYNE function in the phloem vasculature. EMBO Rep 18, 1367–1381.

Ito, Y., Nakanomyo, I., Motose, H., Iwamoto, K., Sawa, S., Dohmae, N. and Fukuda, H. (2006). Dodeca-CLE peptides as suppressors of plant stem cell differentiation. Science 313, 842–845.

Kang, Y. H., Breda, A. and Hardtke, C. S. (2017). Brassinosteroid signaling directs formative cell divisions and protophloem differentiation in Arabidopsis root meristems. Development 144, 272–280.

Khan, J. A., Wang, Q., Sjolund, R. D., Schulz, A. and Thompson, G. A. (2007). An early nodulin-like protein accumulates in the sieve element plasma membrane of Arabidopsis. Plant Physiol 143, 1576–1589.

Kim, H., Zhou, J., Kumar, D., Jang, G., Ryu, K. H., Sebastian, J., Miyashima, S., Helariutta, Y. and Lee, J. Y. (2020). SHORTROOT-Mediated Intercellular Signals Coordinate Phloem Development in Arabidopsis Roots. Plant Cell 32, 1519–1535.

Kinoshita, A., Nakamura, Y., Sasaki, E., Kyozuka, J., Fukuda, H. and Sawa, S. (2007). Gain-of-function phenotypes of chemically synthetic CLAVATA3/ESR-related (CLE) peptides in Arabidopsis thaliana and Oryza sativa. Plant Cell Physiol 48, 1821–1825.

Knoblauch, M., Knoblauch, J., Mullendore, D. L., Savage, J. A., Babst, B. A., Beecher, S. D., Dodgen, A. C., Jensen, K. H. and Holbrook, N. M. (2016). Testing the Munch hypothesis of long distance phloem transport in plants. Elife 5.

Kondo, Y., Nurani, A. M., Saito, C., Ichihashi, Y., Saito, M., Yamazaki, K., Mitsuda, N., Ohme-Takagi, M. and Fukuda, H. (2016). Vascular Cell Induction Culture System Using Arabidopsis Leaves (VISUAL) Reveals the Sequential Differentiation of Sieve Element-Like Cells. Plant Cell 28, 1250–1262.

Kurihara, D., Mizuta, Y., Sato, Y. and Higashiyama, T. (2015). ClearSee: a rapid optical clearing reagent for whole-plant fluorescence imaging. Development 142, 4168–4179.

Lopez-Salmeron, V., Cho, H., Tonn, N. and Greb, T. (2019). The Phloem as a Mediator of Plant Growth Plasticity. Current biology : CB 29, R173–R181.

Lucas, W. J., Groover, A., Lichtenberger, R., Furuta, K., Yadav, S. R., Helariutta, Y., He, X. Q., Fukuda, H., Kang, J., Brady, S. M., et al. (2013). The plant vascular system: evolution, development and functions. Journal of integrative plant biology 55, 294–388.

McBryde, M. C. (1936). A method of demonstrating rust hyphae and haustoria in unsectioned leaf tissue. Am J Bot 23, 686–689.

Musielak, T. J., Slane, D., Liebig, C. and Bayer, M. (2016). A Versatile Optical Clearing Protocol for Deep Tissue Imaging of Fluorescent Proteins in Arabidopsis thaliana. PloS one 11, e0161107.

Pratt, R. B. and Jacobsen, A. L. (2017). Conflicting demands on angiosperm xylem: Tradeoffs among storage, transport and biomechanics. Plant Cell Environ 40, 897–913.

Ren, S. C., Song, X. F., Chen, W. Q., Lu, R., Lucas, W. J. and Liu, C. M. (2019). CLE25 peptide regulates phloem initiation in Arabidopsis through a CLERK-CLV2 receptor complex. Journal of integrative plant biology.

Rodriguez-Villalon, A., Gujas, B., Kang, Y. H., Breda, A. S., Cattaneo, P., Depuydt, S. and Hardtke, C. S. (2014). Molecular genetic framework for protophloem formation. Proc Natl Acad Sci U S A 111, 11551–11556.

Rodriguez-Villalon, A., Gujas, B., van Wijk, R., Munnik, T. and Hardtke, C. S. (2015). Primary root protophloem differentiation requires balanced phosphatidylinositol-4,5-biphosphate levels and systemically affects root branching. Development 142, 1437–1446.

Ross-Elliott, T. J., Jensen, K. H., Haaning, K. S., Wager, B. M., Knoblauch, J., Howell, A. H., Mullendore, D. L., Monteith, A. G., Paultre, D., Yan, D., et al. (2017). Phloem unloading in Arabidopsis roots is convective and regulated by the phloem-pole pericycle. Elife 6.

Ruiz Sola, M. A., Coiro, M., Crivelli, S., Zeeman, S. C., Schmidt Kjolner Hansen, S. and Truernit, E. (2017). OCTOPUS-LIKE 2, a novel player in Arabidopsis root and vascular development, reveals a key role for OCTOPUS family genes in root metaphloem sieve tube differentiation. New Phytol 216, 1191–1204.

Sankar, M., Nieminen, K., Ragni, L., Xenarios, I. and Hardtke, C. S. (2014). Automated quantitative histology reveals vascular morphodynamics during Arabidopsis hypocotyl secondary growth. Elife 3, e01567.

Sanmartin, M., Sauer, M., Munoz, A., Zouhar, J., Ordonez, A., van de Ven, W. T., Caro, E., de la Paz Sanchez, M., Raikhel, N. V., Gutierrez, C., et al. (2011). A molecular switch for initiating cell differentiation in Arabidopsis. Current biology : CB 21, 999–1008.

Smetana, O., Makila, R., Lyu, M., Amiryousefi, A., Sanchez Rodriguez, F., Wu, M. F., Sole-Gil, A., Leal Gavarron, M., Siligato, R., Miyashima, S., et al. (2019). High levels of auxin signalling define the stem-cell organizer of the vascular cambium. Nature 565, 485–489.

Truernit, E. (2014). Phloem imaging. J Exp Bot 65, 1681–1688.

Truernit, E., Bauby, H., Belcram, K., Barthelemy, J. and Palauqui, J. C. (2012). OCTOPUS, a polarly localised membrane-associated protein, regulates phloem differentiation entry in Arabidopsis thaliana. Development 139, 1306–1315.

Truernit, E., Bauby, H., Dubreucq, B., Grandjean, O., Runions, J., Barthelemy, J. and Palauqui, J. C. (2008). High-resolution whole-mount imaging of three-dimensional tissue organization and gene expression enables the study of Phloem development and structure in Arabidopsis. Plant Cell 20, 1494–1503.

Ursache, R., Andersen, T. G., Marhavy, P. and Geldner, N. (2018). A protocol for combining fluorescent proteins with histological stains for diverse cell wall components. Plant J 93, 399–412.

Wendrich, J. R., Yang, B., Vandamme, N., Verstaen, K., Smet, W., Van de Velde, C., Minne, M., Wybouw, B., Mor, E., Arents, H. E., et al. (2020). Vascular transcription factors guide plant epidermal responses to limiting phosphate conditions. Science.

Wilson-Sanchez, D., Martinez-Lopez, S., Navarro-Cartagena, S., Jover-Gil, S. and Micol, J. L. (2018). Members of the DEAL subfamily of the DUF1218 gene family are required for bilateral symmetry but not for dorsoventrality in Arabidopsis leaves. New Phytol 217, 1307–1321.

Zhang, C. and Turgeon, R. (2018). Mechanisms of phloem loading. Curr Opin Plant Biol 43, 71–75.

Zhao, C., Craig, J. C., Petzold, H. E., Dickerman, A. W. and Beers, E. P. (2005). The xylem and phloem transcriptomes from secondary tissues of the Arabidopsis root-hypocotyl. Plant Physiol 138, 803–818.

